# Phasing single-molecule nano-NOMe-seq reveals chromatin state heterogeneity in the context of transcription and long-range interactions

**DOI:** 10.1101/2025.09.08.674887

**Authors:** Stephanie Bellini, Catherine Do, Jane A Skok

## Abstract

A central challenge in molecular biology is determining how 3D chromatin architecture, particularly enhancer-promoter looping and insulating CTCF-mediated interactions, influences gene transcription in individual cells, which has significant implications for healthy and diseased states. To overcome current limitations in imaging and genomic technologies, we developed a cluster-based phasing strategy using long read nano-NOMe-seq to link distinct CTCF binding states— captured at the single molecule level—to the transcriptional status of genes. By stitching partially overlapping long reads and clustering them by shared GpC-accessibility patterns, we stratify CTCF into graded binding states on individual molecules, classify RNA polymerase states at promoters/gene bodies, and infer when spatially separated loci are coordinately activated and occupy loop-competent configurations on the same molecules. When applied to *Sox2, Hoxa*, and *Klf1* regions, cluster-based nano-NOMe-seq phasing reveals how specific topologies bias polymerase behavior and multi-locus activity in ways that bulk assays or locus-engineered imaging cannot fully capture.

## INTRODUCTION

Three-dimensional (3D) organization of chromatin is central to gene regulation. Bulk chromosome conformation capture approaches such as Hi-C and Micro-C generate genome-wide contact maps that reveal two nested axes of organization. First, chromatin segregates into higher-order active (A) and inactive (B) compartments ^3^. Second, within and between these compartments, CTCF and cohesin extrude and stabilize loops that partition the genome into highly self-interactive topologically associating domains (TADs) of ∼1 Mb or less ^4-6^. TADs act as functional units that insulate enhancer–promoter activity, thereby constraining gene regulation within defined boundaries, while this hierarchical architecture brings distant elements into proximity to enable long-range enhancer–promoter interactions that control transcription ^7^.

Beyond these domains, finer-scale interactions within microcompartments were uncovered by Region Capture Micro-C (RCMC) ^8^, that constitute highly focal contacts linking active enhancers and promoters. These interactions persist after acute cohesin depletion or short-term transcriptional blockade by triptolide, indicating that microcompartments do not require loop extrusion or ongoing transcription on short timescales ^8^. However, with extended RNAPII depletion or chronic inhibition, enhancer–promoter contacts weaken globally, consistent with transcription contributing indirectly to the long-term maintenance of these interactions ^9^.

More recently, single-cell Hi-C and related technologies have extended these observations to the level of individual nuclei ^10^. By capturing contact maps in single cells, these approaches can quantify the probability of specific loops or compartmental interactions, though they cannot yet determine whether loop formation and gene expression are coordinated within the same cell. This highlights a key limitation in current technologies: resolving chromatin into its component, heterogeneous states, which reflect the dynamics of interactions in the context of gene activation remains a largely unresolved issue. Indeed, conformation and expression profiling approaches, whether in bulk or single-cell, are uncoupled such that although they can reveal which chromatin contacts are possible or prevalent, and which genes are expressed in a population, they are unable to determine whether active transcription and looping co-occur, or are independent of one another in the same cell for a given locus, and what proportion of alleles/cells are both looped and actively transcribing within a sample.

Live-cell imaging approaches that simultaneously label chromatin and nascent RNA can address this issue. CRISPR/dCas9-based genomic tagging of non-repetitive loci ^11^ has been enhanced by signal-amplifying scaffold systems such as Casilio and CRISPR-Sirius ^12^, which make it possible to visualize weakly fluorescent, low-copy regions in living cells. These chromatin labels can be combined with RNA reporters, such as MS2 or PP7 stem-loop systems that tag nascent transcripts ^13,14^ or more recently dCas13-based RNA imaging strategies. Using this integrated approach, Alexander *et al*., ^15^ showed that transcriptional activation at the *Sox2* locus can occur without stable enhancer–promoter juxtaposition. More recently, Kim *et al*. ^16^ introduced SiCLAT, a dual-color system that simultaneously labels loop anchors and nascent RNA in primary cells, extending live imaging beyond immortalized cell lines. Despite these advances, such techniques remain restricted to a handful of engineered loci.

Complementary fixed-cell imaging methods have provided a powerful means to interrogate chromatin architecture at high resolution in single cells. Optical Reconstruction of Chromatin Architecture (ORCA ^17^) uses sequential oligo-FISH to trace chromatin folding with ∼2 kb resolution across hundreds of kilobases. Applied to the *Sox2* locus, ORCA revealed fine-scale heterogeneous rosette conformations in embryonic stem and neural progenitor cells ^1^. Hi-M ^18,19^ in contrast, was specifically designed to integrate sequential DNA FISH with RNA FISH in intact *Drosophila* embryos, enabling simultaneous visualization of chromatin architecture and transcriptional output at the same loci. Together, these approaches highlight how fixed-cell imaging can achieve either high-resolution structural reconstruction (ORCA) or joint analysis of structure and transcription (Hi-M), though both remain restricted to targeted loci rather than genome-wide views.

Emerging sequencing-based multi-omic single-cell platforms now enable genome-wide coupling of chromatin structure with transcription. Simultaneous single-cell Hi-C and RNA-seq ^20^ directly link genome-wide contact maps with full transcriptomes from the same cells, revealing context-dependent coupling between enhancer–promoter interactions and gene expression. Other strategies, including HiRES ^20^, LiMCA ^20^, and GAGE-seq ^21^, employ variations in library preparation and indexing to balance sensitivity and throughput. These technologies overcome the locus restriction of imaging but remain limited by high sequencing depth requirements and cost. Together, these imaging and sequencing approaches highlight both the progress and current limitations in resolving the temporal and functional coupling of chromatin looping and transcription at single-cell resolution.

To overcome these constraints, we made use of long read nano-NOMe-seq to link distinct CTCF binding states—captured at the single molecule level—to the transcriptional status of genes. Using a cluster-based approach analogous to pseudobulk analyses in single-cell datasets, we could grade CTCF binding at a near single cell level, classify Pol II states at promoters and across gene bodies, and infer when dispersed loci are co-activated and in loop-ready configurations on the same molecules. Applied to the *Sox2, Hoxa*, and *Klf1* regions, this cluster-phased view reveals how specific topologies bias polymerase behavior and multi-locus activity—features not fully captured by ensemble assays or engineered-locus imaging.

## RESULTS

### Clustering of nucleosome occupancy reveals graded CTCF binding and Pol II–dependent transcription states

To identify distinct within-sample binding states for CTCF and Pol II, we performed long-read Nano-NOMe-seq in mESCs. This enabled us to assess genome-wide nucleosome positioning using exogenous 5mC at HGC (GC excluding CGC) at the single molecule (SM) level^22^ (**Figure 1A**). Unlike MNA-seq analyses, the single molecule (SM) data can be clustered to reveal different CTCF and RNA Pol II binding states based on nucleosome occupancy patterns. For CTCF, we analyzed a 2 kb region centered on CTCF binding sites (CBSs) validated by ChIP-seq. As shown in the heatmap, this uncovered a continuum of binding states previously described by us ^22^ (**Figure 1B**). Since nano-NOMe-seq libraries are PCR-free, for a given locus, the proportion of each SM cluster is likely to reflect the percentage of cells with similar CTCF binding states.

**Figure 1.**
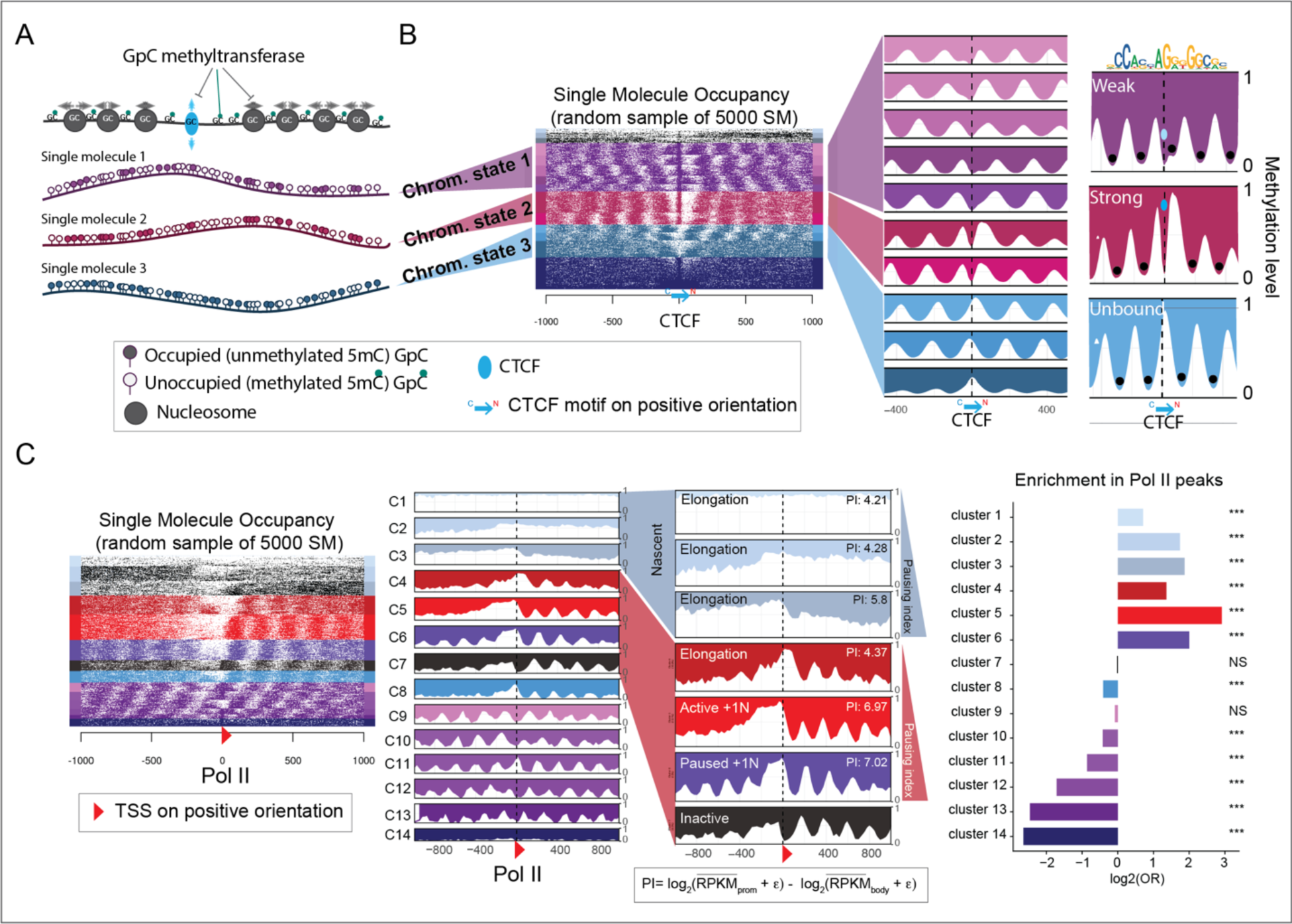
Clustering of nucleosome occupancy patterns from Nano-NOMe-seq reveals graded CTCF binding and Pol II–dependent transcription states. **A**. Cartoon illustrating the Nano-NOMe-seq strategy to identify single-molecule (SM) nucleosome positioning and transcription factor (TF) footprinting, based on exogenous 5mC levels in a GpC context. **B**. Heatmap and average profiles of 2 kb regions centered on CTCF binding sites, showing clustering of SMs in mESCs to define within-sample CTCF binding states. **C**. Heatmap and average profiles of 2 kb regions centered on Pol II–bound transcription start sites (TSSs), showing clustering of SMs in mESCs to define within-sample Pol II transcription states. Hyper-accessible clusters, C1-C3, were color-coded in blue and annotated as nascent chromatin ^2^. Clusters are ordered and annotated based on Pol II pausing index (PI) and enrichment for Pol II peaks. ε=0.5.

To classify CTCF binding states, we aggregated clusters as unbound (light blue – no CTCF footprint), weak/transitioning (purple – CTCF footprint adjacent or overlapping a nucleosome) or strong (red – CTCF footprint within a nucleosome-free region-NFR). A similar analysis was performed to identify within-sample Pol II transcription states, centering on ChIP-seq validated, Pol II–bound transcription start sites (TSSs). SM clustering in the heatmap was ordered and annotated based on a Pol II pausing index (PI) and enrichment for Pol II peaks (**Figure 1C**). Using this approach, we classified sites as elongating, active with +1 nucleosome, paused and inactive.

### Clustered-based phasing of Nano-NOMe-seq single molecules

The goal of these studies was to link distinct CTCF binding states—captured at the single molecule level—to the chromatin and transcriptional status of genes and their regulatory elements, however the median read length of the SM nano-NOMe-seq was not sufficient for this purpose (**Figure 2A** left) and stitching reads together was necessary to be able to incorporate CTCF loop bases with intervening regulatory elements and genes across a region of interest. **Figure 2A (**right) shows an example of SM methylation sequence at the GpC level across a locus spanning 25 kb.

**Figure 2.**
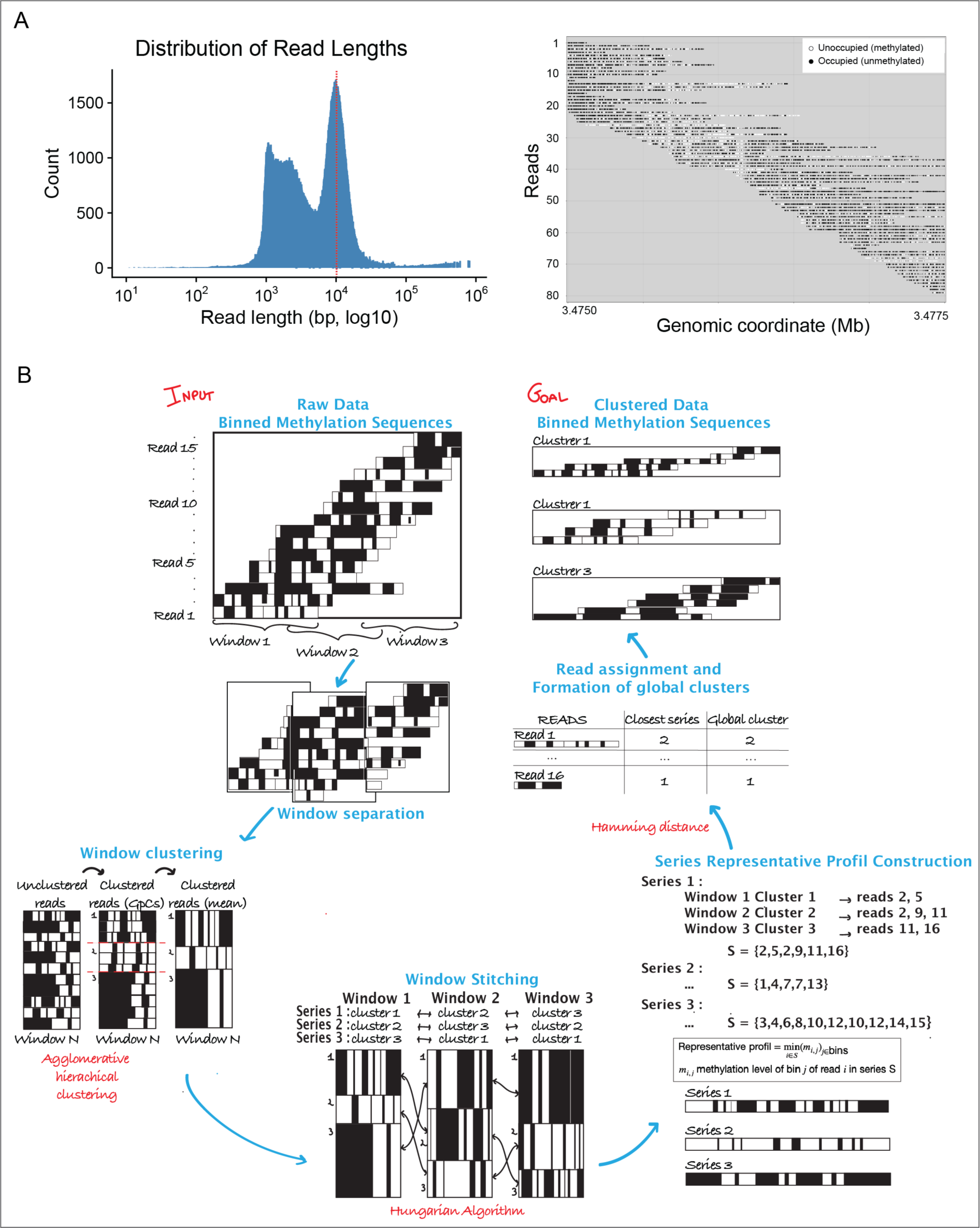
Clustered-based phasing of the Nano-NOMe-seq single molecules. **A**: Left, distribution of Nano-NOMe seq read lengths. The read dashed line indicates the median. Right, example of SM methylation sequence at the GpC level in a locus spanning 25 kb. **B**: For a given long-range locus, single molecules (Nano-NOMe-seq reads) are binned into 50 bp. The region is divided into overlapping windows to enable stitching: SMs are first clustered within each window using agglomerative hierarchical clustering, and clusters are then matched across consecutive overlapping windows using the Hungarian algorithm. After generating aggregated methylation profiles per cluster, each SM is reassigned to its best-matched cluster based on Hamming distance. This step corrects inconsistent assignments across windows and yields long-range phased SM clusters, which can be analyzed at either the bin or GpC level.

Our approach for stitching is described in the scheme in **Figure 2B**. Nano-NOMe-seq reads from a given long-range locus were binned into 50 bps, to obtain sufficient resolution to detect both transcription factor and nucleosome footprints. The locus was then divided into overlapping windows to facilitate stitching. SMs were clustered within each window using agglomerative hierarchical clustering (**Figure S1A**), and clusters matched across consecutive overlapping windows using the Hungarian algorithm (**Figure S1B**). After generating aggregated methylation profiles for each cluster, SMs were reassigned to their best-matched neighboring clusters based on the Hamming distance (**Figure S1C**). This step corrects inconsistent assignments across windows and yields long-range phased SM clusters, which can be analyzed at either the bin or GpC level.

An example of cluster-based phasing for a locus spanning 50 kb is shown in **Figure 3A**, with a cartoon illustrating the approach used to estimate the accuracy of the algorithm across 90 extended loci, each mapped by at least one long (≥50 kb) read that is kept out of the analysis and used for validation (**Figure 3B** and **Table S1**). For each locus, the methylation sequence of the long read is compared to the mean methylation sequences of all clusters (c_j_), and the closest cluster is identified using the Hamming distance (d_H_). A(r_i_) denotes the read accuracy, reflecting how well read r_i_ matches its best-aligned cluster (higher values = better match). The overall clustering accuracy is defined as the average of A(r_i_) across all long reads. The distribution of read accuracy across all long reads is shown in **Figure 3C**. The global clustering accuracy was 0.78. Examples of long-read methylation sequences and their closest-matching clusters for three loci spanning 50–60 kb are shown in **Figure 3D**.

**Figure 3.**
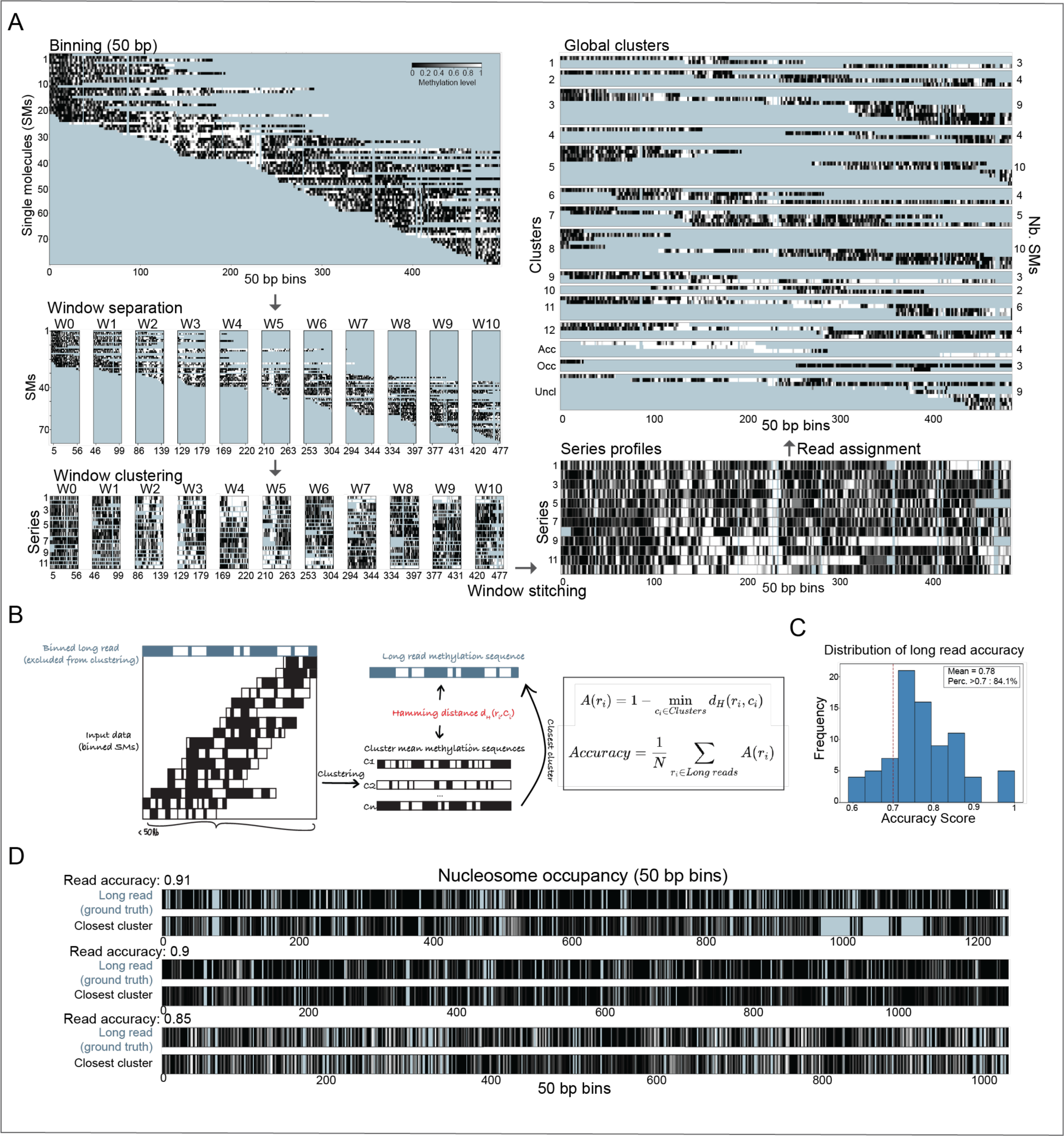
Examples and accuracy of Nano-NOMe-seq single molecules clustered-based phasing. **A**: Example of the clustered-based phasing algorithm for one locus spanning 50 kb. Missing data are color-coded in blue. **B**: Cartoon illustrating the approach used to estimate the accuracy of cluster-based phasing across 90 long-range reference loci, each mapped by at least one long (≥50 kb). For each locus, the methylation sequence of a long read is compared to the mean methylation sequences of all clusters (c_j_), and the closest cluster is identified using the Hamming distance (d_H_). A(r_i_) denotes the *read accuracy*, reflecting how well read r_i_ matches its best-aligned cluster (higher values = better match). The overall clustering accuracy is defined as the average of A(r_i_) across all long reads. **C**: Distribution of read accuracies across all long reads. The global clustering accuracy is 0.78. **D**: Examples of long-read methylation sequences and their closest-matching clusters for three loci spanning 50–60 kb.

### Cluster-based phasing of nano-NOMe-seq data confirms CTCF-independent Sox2 transcription

As proof of principle, cluster-based stitching/phasing of nano-NOMe-seq data was implemented at the well-studied *Sox2* locus to determine whether there is regulatory heterogeneity in mESCs. *Sox2* expression is regulated by the *Sox2* control region. The SCR contains two distal enhancers, SRR107 and SRR111, which flank a CTCF binding site known as SRR109. Previous studies have shown that, while deletion of the SCR in mice significantly reduces Sox2 expression ^23^, deletion of the CTCF motif at SRR109 results in only a modest, non-significant decrease in Sox2 levels ^24 25^, suggesting that transcription is primarily mediated by CTCF-independent enhancer–promoter interactions ^26^.

To determine whether we could confirm these results, we performed cluster-based phasing of SMs using our previously generated nano-NOMe-seq data ^22^. The average coverage and median read lengths mapping the *Sox2* locus are reported in **Table S2**. Region Capture Micro-C (RCMC) ^8^ (at 200 bp resolution) of the 150 kb *Sox2* locus in mESCs revealed a previously described CTCF-mediated loop between the *Sox2* control region (SCR) and a CTCF binding site (window 1) located 2 kb upstream of the *Sox2* transcription start site (TSS, window 2) (**Figure 4A**). The two SCR enhancers, SRR107 (window 4) and SRR111 (window 6), flanking the downstream CTCF anchor (SRR109, window 5) are highlighted. CTCF and Pol II ChIP-seq (shown in **Figure 4A**) confirmed binding of CTCF and active Sox2 transcription / elongation respectively in mESCs.

**Figure 4:**
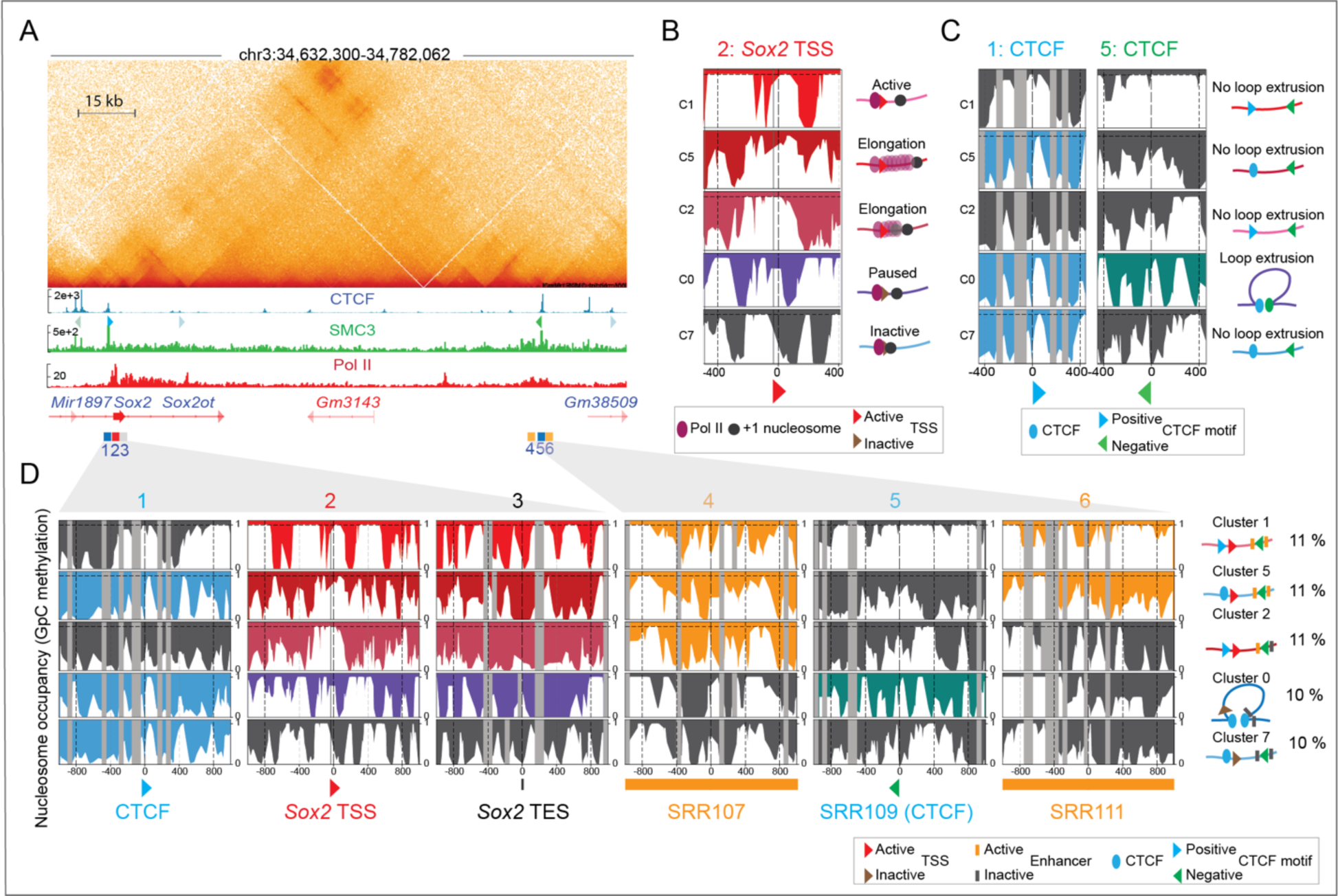
Cluster-based phasing of nano-NOMe-seq data confirms CTCF-independent Sox2 transcription. **A**: Capture Micro-C (300 bp resolution) of the 150 kb Sox2 locus in mESCs (GSE207225) reveals a CTCF-mediated loop between the Sox2 control region (SCR) and a CTCF binding site (window 1) located 2 kb upstream of the Sox2 transcription start site (TSS, window 2). The SCR includes two enhancers, SRR107 (window 4) and SRR111 (window 6), flanking the downstream CTCF anchor (SRR109, window 5). CTCF motif orientations are shown below the CTCF ChIP-seq track. Pol II ChIP-seq (GSE207225) confirms active Sox2 transcription and elongation in mESCs. **B:** Single-molecule nucleosome occupancy clustering identifies distinct transcriptional states within the sample, reflecting dynamic Pol II pausing, release, and elongation. This window is centered on Pol II with the TSS located 30 bp downstream. **C:** Nano-NOMe-seq at CTCF anchors shows that only a subset of molecules exhibits a clear CTCF footprint and associated nucleosome-free region (NFR) at both anchors. **D:** Cluster-based phasing resolves the coupling between loop extrusion, enhancer–promoter interactions, and Sox2 transcription. Clusters 1, 2 and 5 exhibit CTCF-independent Sox2 transcription, whereas cluster 0 links Sox2 Pol II paused transcription to CTCF-mediated loop extrusion. SRR107 is accessible (active) across all clusters with active Sox2 transcription, while SRR111 is inactive in Cluster 2, where elongation is reduced. In cluster 0 and cluster 7, both SRR107 and SRR111 are inaccessible (inactive). CTCF motifs are colored according to orientation: blue for positive and green for negative.

In this locus, our cluster-based phasing approach allowed us to resolve regulatory elements within 9 clusters of single molecules (**Figure S2**). Clusters 0, 1, 2, 5, 7 displayed nucleosome patterns indicative of distinct transcriptional states within the sample, reflecting dynamic Pol II activation with +1 nucleosome, elongation and pausing, (**Figure 4B)**. Nano-NOMe-seq at CTCF anchors shows that only a subset of molecules exhibited a clear CTCF footprint and associated nucleosome-free region (NFR) at both anchors (**Figure 4C**). We hypothesized that clusters exhibiting CTCF footprint at chromatin loop anchors identified by chromatin capture assays and overlapping CTCF motifs arranged in a convergent orientation (blue – forward, green – reverse), likely represent the subset of cells harboring a CTCF-mediated loop. At the *Sox2* locus, only ∼10% of the SMs/cells harbored a CTCF-mediated loop between the SCR and the CTCF binding site located directly upstream of *Sox2* TSS, confirming the high within-sample heterogeneity of chromatin architecture and CTCF binding states.

**Figure 4D** shows the relationship between *Sox2* transcription, enhancer activity at SRR107 and SRR111, and CTCF-mediated looping, resolved at the SM cluster level. Clusters 1, 2 and 5 exhibit CTCF-independent *Sox2* transcription (no CTCF footprints present), whereas cluster 0 links *Sox2* Pol II paused transcription to the presence of CTCF footprints that could mediate the formation of loops detected by RCMC. SRR107 is accessible (active) across all clusters with active *Sox2* transcription, while SRR111 is inactive in cluster 2, where elongation is reduced. In cluster 0 and cluster 7, both SRR107 and SRR111 are inaccessible (inactive). These results validate previous findings ^15,24-26^, and explain why deletion of the CTCF motif in this region does not impact *Sox2* expression. Furthermore, they highlight the connections between SRR107 and SRR111 activity and elongation strength: elongation is more pronounced when both enhancers are accessible and active, and reduced when only SRR107 is active.

### Cluster-based phasing identifies CTCF footprints at the bases of long-range multiway loops including the *Sox2* locus

We next extended our analyses downstream of the *Sox2* locus and SCR, to examine CTCF footprints at bases of CTCF-mediated rosettes identified by ORCA ^1^ and loops also shown to be present in RCMC ^8^ (**Figure 5A** and **Table S2**). Because of the reduced coverage within overlapping windows over the extended region, we identified fewer clusters (7 in total, **Figure S3**). We detected CTCF footprints and binding sites in the appropriate orientation to form loops in clusters 1, 2 and 5 (arcs in **Figure 5A and 5B**). Our cluster-based analysis confirmed the presence of CTCF footprints at CTCF anchors of the *Sox2* rosette (windows 1, 5 and 6) previously validated by ORCA ^1^, but revealed a relatively weak CTCF footprint in window 6. Based on the strength of the CTCF footprint and the finding that strong footprints highlight strong CTCF signals that are better able to stop cohesin ^27^, we expect that stronger loops form between CBS sites with stronger CTCF footprints (solid lines) and weaker loops form between CBS sites with weaker CTCF footprints (dashed lines). Interestingly, we identified two CTCF footprints that could mediate long-range loops connecting window 1 with windows 9 and 10 in cluster 5, and window 1 with window 10 in cluster 6 that can be visualized by RCMC, but that were not detected with ORCA. These findings suggest that loops detected by bulk interaction data in the *Sox2* locus arise from a mixture of two-way and higher-order chromatin interactions consistent with distinct rosette formations (**Figure 5B**).

**Figure 5:**
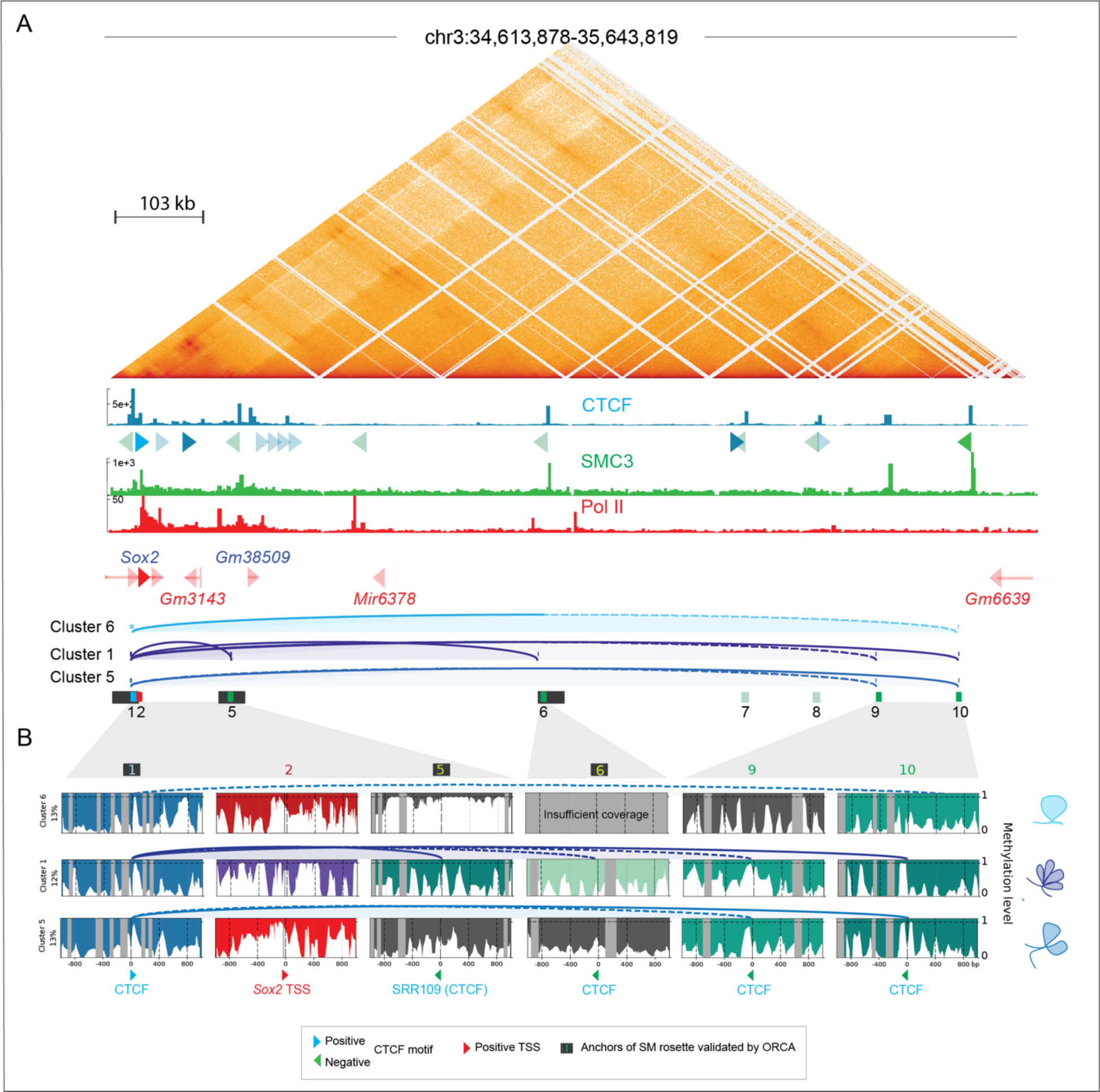
Cluster-based phasing of nano-NOMe-seq data identifies CTCF footprints at the bases of long-range multiway loops surrounding the *Sox2* locus. **A:** Capture Micro-C (1 kp resolution), CTCF, SMC3 and Pol II ChIP-seq across the extended 1 Mb *Sox2* locus in mESCs. Arcs depict CTCF-dependent loops predicted by nucleosome and CTCF footprinting in clusters with at least one predicted loop. The black rectangles highlight CTCF anchors of the *Sox2* rosette previously validated by ORCA ^1^. **B**: Nano-NOMe-seq profiles of the 2 kb windows centered on the CTCF motifs involved in looping and on *Sox2* TSS. Arcs represent predicted CTCF-dependent loops with dashed line corresponding to loops connecting at least one anchor with a weaker CTCF footprint.

We did not observe an association between predicted *Sox2* transcription and higher-order CTCF-dependent interactions. Consistent with our findings in **Figure 3**, both cluster 5 and cluster 6 chromatin conformations were permissive for *Sox2* expression in the absence of a potential contact between the CTCF site at the TSS and the SCR (SRR109). In contrast, co-occurring multiway loops between the downstream and both upstream CTCF sites (cluster 5) were linked to weaker Pol II elongation. Since deletion of CTCF at the downstream sites has no effect on Sox2 expression ^26^, we conclude that the multiway loops we detect do not contribute to SCR-mediated effects on transcription in ESCs. However, we cannot rule out that in other contexts these may be important for Sox2 gene regulation. Together, our cluster-based nano-NOMe-seq CTCF and Pol II footprint analysis highlight the potential for identifying rosette structures across extended regions, and the connections between loop formation and Pol II status.

### Cluster-based phasing of nano-NOMe-seq identifies long-range high-order multiway CTCF dependent loops across the *Hoxa* region

CTCF-mediated rosette structures were also previously identified by ORCA across the *Hoxa* region ^1^. Using our cluster-based nano-NOMe-seq we asked if we could identify nucleosome and CTCF footprinting at the bases of these loops, together with the concomitant Pol II occupancy state (**Table S2**). Our previously generated Hi-C data ^27^ revealed pairwise chromatin interactions between ORCA-identified loop anchors and additional CTCF/SMC3 sites across the 2.3 Mb domain (**Figure 6A)**. ORCA loop anchors corresponded exclusively to bound CTCF motifs in the negative orientation, suggesting that additional upstream CTCF binding sites in the positive orientation contribute to the rosette structure identified by imaging analysis. Among the nano-NOMe-seq SM clusters, only cluster 5 exhibited co-occurring CTCF footprints at all three ORCA loop anchors (**Figure 6B**). In this cluster, we also identified two weak upstream CTCF footprints (window 1 and 2) mapping to motifs in the positive orientation that could enable the rosette formation by providing the required convergent CTCF binding. These were not included in the ORCA analyses of three-way interactions. Cluster 5 additionally showed footprints at multiple CTCF binding sites in both orientations between window 3 and 9, consistent with higher-order multiway interactions beyond those originally described. The other clusters displayed a spectrum of two-way and multiway interactions involving distinct CTCF binding sites within the domain (**Figure 6B**), again underscoring chromatin conformation heterogeneity not captured by bulk assays and a complexity of high-order interactions that might not be fully anticipated in SM targeted designs.

**Figure 6:**
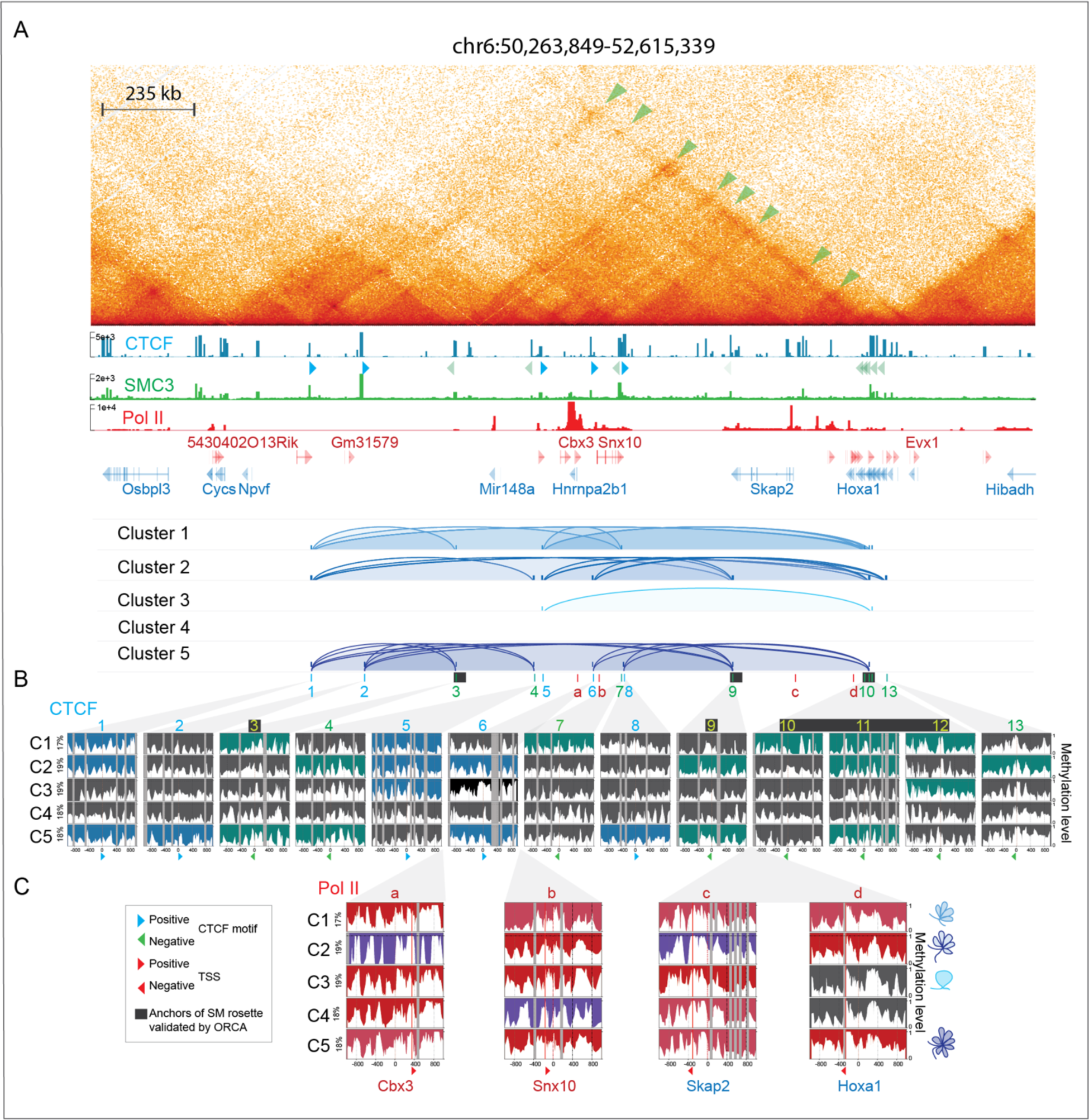
Cluster-based phasing of nano-NOMe-seq identifies long range high-order multiway CTCF dependent loops across the *Hoxa* region. **A:** Hi-C (5 kb resolution), CTCF, SMC3 and Pol II ChIP-seq across the 2.3 Mb *Hoxa1* locus in mESCs. Arcs show CTCF-dependent loops predicted by nucleosome and CTCF footprinting in each cluster. The black rectangles show CTCF anchors of the *Hoxa1* rosette previously validated by ORCA ^1^. **B**: Nano-NOMe-seq profiles of the 2 kb windows centered on the CTCF motifs overlapping with the anchors of loops identified by Hi-C and ORCA. Sites with CTCF footprints and their potential for loop formation are highlighted by the orientation of the CTCF motif (blue – forward, green – reverse). **C**: Nano-NOMe-seq profiles of the 2 kb windows centered on Pol II summits at the TSS of genes within CTCF loops, colored according to Pol II states defined in **Figures 1** and **4**. The cartons on the right represent the chromatin conformation predicted by CTCF footprint (shown in **B**) for each cluster.

While ORCA has identified the presence of rosettes within extended regions, it does not determine whether these support or inhibit gene expression. In contrast, our nano-NOMe-seq cluster-based approach has the potential to predict within-sample looping associated with Pol II activity. The Pol II activation status across the *Hoxa* region in each rosette-forming cluster, demonstrates that some rosette structures are compatible with activity across the entire region, while others are linked to Pol II pausing and inactivity at particular sites (**Figure 6C**). As a general rule, we observe that higher connectivity across the *Hoxa* region is associated with higher Pol II activity.

### Cluster-based phasing of nano-NOMe-seq data reveals long-range high-order multiway CTCF independent micro-compartments loops across the *Klf1* region

We next turned our attention to the *Klf1* region, which RCMC identified as being part of a microcompartment with multiway interactions (**Figure 7A** and **Table S2**). Although microcompartments are not (at least in the short term) reliant on either loop extrusion or transcription, contacts largely occurred between active enhancers and promoters that are enriched for Pol II, as highlighted by RCMC (**Figure 7A**) and Pol II peaks were overlapping A-micro-compartments (red) as shown by the micro-compartment score (PC2 values from Hi-C eigenvector decomposition).

**Figure 7:**
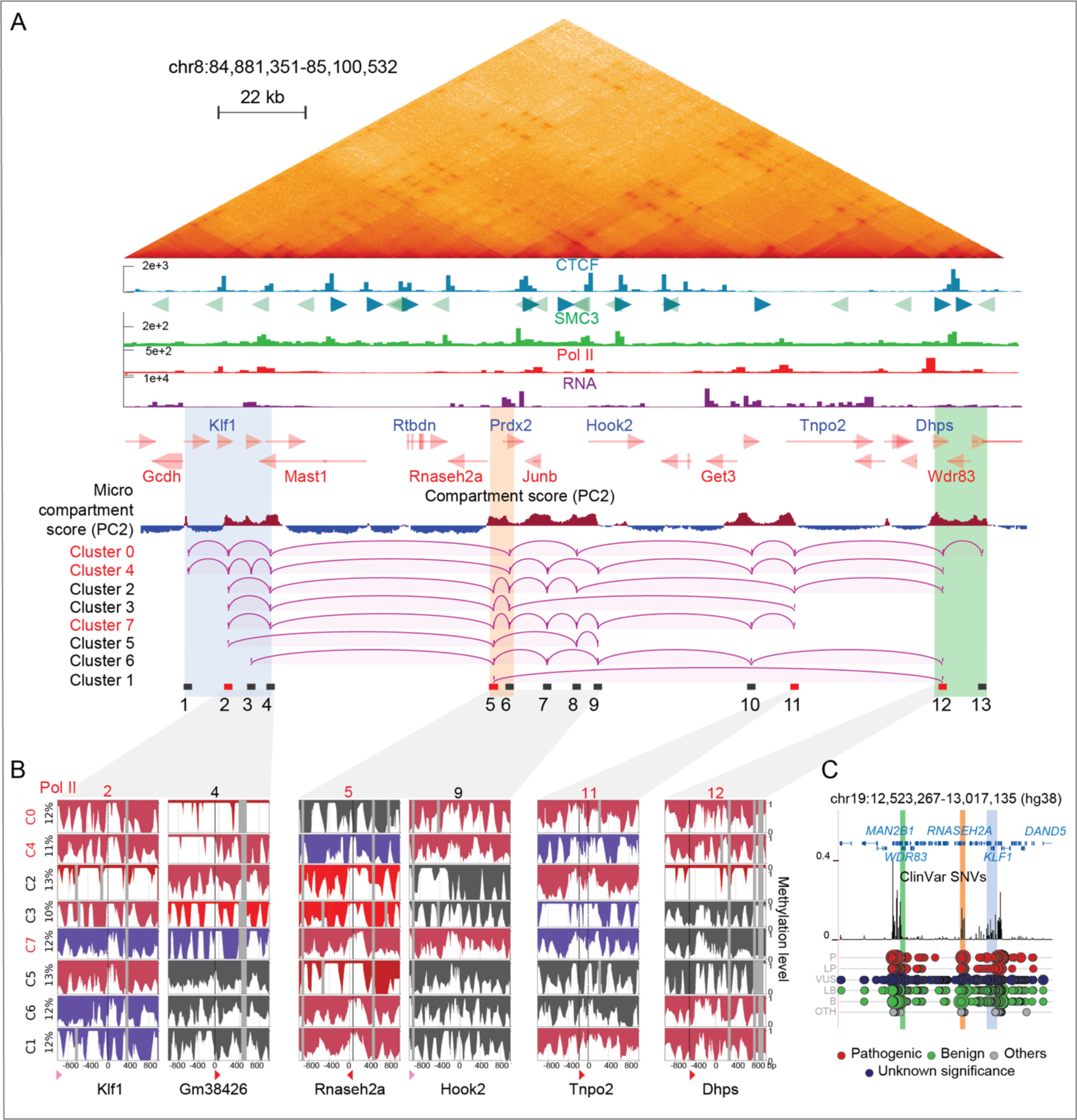
Cluster-based phasing of nano-NOMe-seq data reveals long range high-order multiway CTCF independent micro-compartments loops across the *Klf1* region. **A:** RCMC (300 bp resolution), CTCF, SMC3, Pol II ChIP-seq and RNA-seq across the 220 kb *Klf1* region in mESCs. The micro-compartment score (PC2 values from Hi-C eigenvector decomposition) highlights A-micro-compartments (red), which overlap Pol II peaks. Arcs depict CTCF-independent loops predicted by nucleosome occupancy at Pol II sites in each cluster. Red rectangles indicate anchors with predicted interactions in ≥ 5 out of 8 clusters. Clusters supporting the highest-order multiway interaction are also highlighted in red. **B**: Nano-NOMe-seq profiles of the 2 kb windows centered on Pol II summits overlapping A-micro-compartments involved in looping. **C**: ClinVar single-nucleotide variant (SNV) track in the syntenic human region, showing peaks of pathogenic SNVs overlapping chromatin interaction hubs that connect the center and boundaries of the region. (L)P (likely) Pathogenic; (L)B, (likely) begnin; VUS, variant of unknown significance; OTH, other.

Using our cluster-based nano-NOMe-seq we analyzed the nucleosome and Pol II status across the region, centering our analysis on Pol II peaks. We assumed that TSSs exhibiting co-occurring active Pol II-associated nucleosome patterns likely represent co-activated genes engaged in multiway interactions across A micro-compartments identified by RCMC ^8^. Across 8 windows, we identified clusters with distinct combinations of elongating, paused or inactive Pol II sites (**Figure 7B** and **S4**). Clusters containing multiple simultaneously elongating Pol II sites, predicted by nucleosome occupancy, are shown in **Figure 7B**. Arcs indicate the connections between simultaneously active sites in each cluster, which are highlighted as CTCF-independent contacts in the RCMC map. For visualization, only consecutive predicted interactions are displayed. Although we cannot exclude the existence of additional pairwise combinations within a given cluster, no cluster exhibited a single loop spanning the whole 220 kb region. Cluster 1, representing only 10 % of the SM population, was the only case with a single loop spanning 115 kb, consistent with the reported maximum loop length achievable without CTCF-mediated loop extrusion ^28,29^. Red rectangles in **Figure 7B** indicate windows with coordinated activity in ≥ 5 out of 8 clusters. The coalescence of active sites in these regions suggests coordinated multiway interactions forming activity hubs at three locations: a central region (window 5) bridging two edge regions located beyond the distance where chromatin interactions typically form without CTCF. These hubs encompass genes including *Klf1, RNaseH, Jun2, Tnp2* and *Dhps*, suggesting coordinated regulation of these genes.

Since many of the genes in these activity hubs are important for function and survival, we turned to the ClinVar single-nucleotide variant (SNV) track in the syntenic human region. Intriguingly, peaks of pathogenic SNVs overlap all three major activity hubs, connecting the center and boundaries of the region, highlighting the relevance of the hubs.

## DISCUSSION

Despite advances in long-read sequencing, current technologies such as nanopore remain insufficient to generate ultra-long reads that span entire TADs at the depth required for comprehensive analysis. We generated a single-molecule, cluster-based phasing strategy for nano-NOMe-seq (analogous to pseudobulk in single-cell data) that resolves heterogeneous chromatin states and their coordination with transcription and long-range CTCF/cohesin architecture. Stitched single-molecule clustering of nano-NOMe-seq delivered near single-cell resolution to: (i) map how discrete Pol II states (elongating, active, paused, inactive) align with CTCF footprints at defined loop anchors—explicitly revealing when dual-anchor occupancy is compatible or incompatible with elongation; (ii) detect simultaneous multi-anchor CTCF occupancy consistent with cohesin-mediated rosette topologies, while on the same molecules, identifying which conformations are permissive for Pol II activation; and (iii) identify co-elongating loci within microcompartments on single molecules, exposing recurrent, coordinated activity hubs. PCR-free nano-NOMe preserves molecule counts, so cluster proportions approximate within-sample allele/cell fractions. This framework not only reproduces prior findings but also adds mechanistic resolution into how topology partitions transcriptional states across complex regulatory domains.

By integrating cluster-level CTCF footprint and nucleosome patterns with ChIP-seq and chromatin capture data, we could predict cluster-specific chromatin interactions and transcriptional activity that were validated at the bulk level, thereby deconvolving distinct chromatin conformations that enforce distinct transcriptional outcomes. The graded spectrum of CTCF footprints (unbound → transitioning → strong NFR-embedded) likely reflects differences in residence time and cohesin-stopping efficiency, providing a mechanistic framework for interpreting how anchor occupancy shapes transcriptional compatibility. At the *Sox2* locus, for example, strong simultaneous CTCF footprints at the promoter-proximal anchor and the SCR-embedded SRR109 anchor (window 5) were selectively enriched on paused molecules but absent in elongating clusters, indicating that the 1↔5 configuration is not compatible with elongation. In contrast, elongation was observed when SRR109 (window 5) lacked a CTCF footprint, or when the promoter-proximal anchor paired with the further distal downstream anchors (windows 9/10). These observations reconcile why deleting the SRR109 motif does not affect steady-state Sox2 expression as enhancer activity drives transcription when the 1↔5 loop cannot form. In mESCs, deletion of CTCF at the promoter-proximal anchor has been shown to have no effect on Sox2 expression ^26^, indicating that the longer distance multiway loops that contribute to rosette formation are dispensable for SCR-mediated transcription in ESCs. However, they may be important for Sox2 gene regulation in other contexts. Consistent with this, the enrichment of paused Pol II in the 1↔5 configuration suggests a dynamic poised state in which Pol II remains primed for rapid activation or repression. Such poised states are frequently involved the regulation of developmental genes ^30^. In this view, the 1↔5 loop may not simply be an architectural feature, but a regulatory checkpoint that modulates transcriptional responsiveness—stabilizing a paused Pol II complex until appropriate contextual signals trigger either productive elongation or silencing. Importantly, even when bulk contact frequencies appear stable, transcriptional changes may reflect a redistribution of single-molecule state ensembles (e.g., looped-paused vs unlooped-elongating) rather than loss of specific contacts. This highlights how specific CTCF-mediated topologies can gate transcriptional plasticity, linking three-dimensional genome organization with dynamic gene expression.

Cluster-based phasing of SMs across hundreds of kilobases enabled us to deconvolve bulk pairwise chromatin interactions into a mixture of two-way and multiway interactions consistent with ‘rosette’ architectures, previously described at the *Sox2* and *Hoxa* domains using SM imaging ^17^. However, the non-targeted design of NOMe-seq data uncovered an unexpectedly rich spectrum of cluster-specific interaction patterns that extended beyond the originally queried regions, allowing unbiased discovery of higher-order conformations and revealing a level of complexity not fully appreciated in earlier assays. Crucially, this molecule-resolved view separates structural compatibility from transcriptional activity. Unlike long-read chromatin capture methods such as Pore-C, which identify multiway interactions at SM resolution ^31^, but cannot resolve their impact on transcription, our approach couples 3D architecture with Pol II and nucleosome status, thereby linking specific conformations to their functional output. For example, we showed that *Hoxa* clusters with higher predicted connectivity were more often permissive for polymerase engagement across the region, but distinct rosette ensembles were compatible with the activation of different gene combinations. Together, these results refine the rosette concept: rather than a single hub, rosettes constitute ensembles of loop-competent configurations, distinguished by which anchors are simultaneously involved. These distinct arrangements bias promoter states toward pausing or activation, depending on local regulatory grammar, anchor strength, and enhancer wiring.

Our nucleosome occupancy-based approach is not limited to predicting CTCF-mediated multiway loops. At the *Klf1* locus, for example, microcompartment organization has previously been identified using bulk RCMC. Although such microcompartment contacts persist after acute inhibition of transcription, our single-molecule data show that their most active instances correspond to concurrent elongation constellations across promoters and enhancers—a coordination uniquely captured by nano-NOMe. Similar to the rosette structures, stitched clusters revealed synchronous elongation across multiple Pol II peaks on the same molecules, generating an even more diverse repertoire of multiway combinations. Despite this complexity, three recurrent activity hubs emerged, coinciding with A-compartment foci. The central hub bridged distal regions that, in the absence of this configuration, would otherwise require CTCF-mediated looping to interact. These findings suggest that such hubs encompass genes requiring coordinated regulation. Supporting this hypothesis, these activity regions overlap with peaks of pathogenic human SNVs in the syntenic interval, suggesting that coordinated microcompartment organization may mark regulatory hotspots particularly vulnerable to disease-associated variation.

As with any predictive framework, our conclusions are restricted by the indirect nature of accessibility footprints, inference of Pol II states, coverage depth, and gaps in GpC density, which together constrain resolution and sensitivity; nevertheless, our approach already overcomes key limitations of existing methods by linking chromatin architecture to transcriptional outcomes at near single-molecule resolution, and future methodological advances will only expand its power.

In conclusion, cluster-based single-molecule phasing from nano-NOMe-seq data reveals that regulatory loci exist as ensembles of loop-competent and transcriptionally distinct states whose proportions—rather than population averages—govern output. Our framework bridges the gap between contact maps and functional engagement and provides a roadmap for perturbations that shift state distributions to modulate gene expression.

## MATERIALS AND METHODS

### Datasets and Data Processing

#### Nano-NOMe-seq

Nano-NOMe-seq data (GSE280711) were previously generated in mESCs and processed as described in Do *et al*.,^22^. Briefly, base modification calling and alignment against the mm10 reference genome were performed using Dorado. GpC methylation of single molecules (SMs) mapping to the regions of interest was extracted using modkit extract.

#### Hi-C

Hi-C data (GSE270669) were previously generated using the Arima Hi-C kit (A510008), aligned to mm10, and processed with HiC-Pro and HiCExplorer, as described in Do *et al*., ^27^. Hi-C matrices were generated at 10 kb resolution.

#### Capture Micro-C

Capture Micro-C interaction matrices (50 bp resolution) aligned to mm39 were downloaded from Goel *et al*., ^8^ (GSE207225). The 2 replicates were merged, zoomified and balanced using cooler.

#### ChIP-seq, RNA-seq, and ATAC-seq

CTCF and SMC3 ChIP-seq, RNA-seq, and ATAC-seq data (GSE270669) were previously generated in mESCs, aligned to mm10, and processed as described in Do *et al*., ^27^, using the Seq-N-Slide pipeline. Pol II ChIP-seq data aligned to mm39 were downloaded from Goel *et al*., ^8^ (GSE207225), and subsequently realigned to mm10 using the Seq-N-Slide pipeline.

#### Visualization

Genomic maps were generated using HiGlass and IGV. For the *Sox2* and *Klf1* regions, ChIP-seq and RNA-seq data were lifted over to mm39 to ensure consistency with the Capture Micro-C data. For the *Klf1* locus, the micro-compartment score was defined as the second component (PC2) eigenvector values derived from eigenvector decomposition of the Hi-C correlation matrix and was generated using hicPCA (HiCExplorer). For the *Sox2* and *Hoxa1* loci, coordinates of anchors of multiway interactions identified by ORCA were obtained from *Hafner et al*., ^1^.

### Nano-NOMe-Seq clustered-based phasing

#### Formatting and Filtering

Preprocessed files were imported and formatted to group GpC methylation calls by read identifier. For each GpC site, the mean coverage was calculated. Reads were then filtered to exclude low-quality or uninformative molecules based on length (<10 bp) and the number of methylated sites retained (<10 GpCs).

#### Binning

Methylation sequences were partitioned into 50 bp bins spanning from the first read start to the last read end. Smoothed methylation matrices were generated across fixed bins. *These matrices enabled* visualization of methylation density profiles and identification of fully methylated versus fully unmethylated reads. Because those reads were less informative to assess footprinting and can be ambiguous for stitching, they were pre-assigned to a fully methylated and unmethylated cluster, respectively and ignored for downstream analyses.

#### Window Separation

To enable localized clustering, sliding windows were defined across the genomic bins, and valid methylation sequences were extracted for each window. Windows were defined as 60 valid bins (valid bin were defined as bin with at least one read with methylation data) with at least 20 overlapping bins. Reads with less than 30 valid bins were excluded.

#### Window-Based Clustering

Within each window, pairwise similarity between methylation sequences was computed using a masked Hamming distance metric that ignores missing values. Agglomerative hierarchical Clustering was then performed within windows, ensuring rare cell states could be captured even if not present across the full region. This algorithm separates the reads within each window into *t* clusters based on the similarity of their methylation patterns: each data point is first treated as an individual cluster. At each iteration, the two closest clusters are merged based on the average linkage (average of all pairwise distances between points in the two clusters). This process continues until the desired number of clusters, *t*, is reached. The agglomerative hierarchical clustering procedure is described in detail below:

Let *W d*enote a window of the dataset containing *n* reads, with corresponding methylation sequence vectors {*x*_1_, *x*_2_, …, *x*_*n*_}.

The clustering algorithm input is a distance matrix *D*, where each element *d*_*H*_(*u, v*) represents the masked Hamming distance between two methylation sequence vectors *u* and *v*. The distance is defined as:

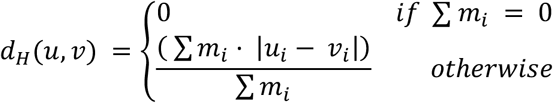

where *m*_*i*_ = 1 if both *u*_*i*_ and *v*_*i*_ are observed (non-missing) and *m*_*i*_ = 0 otherwise.

Step 1. Initialization: Start with the *n* reads in the window, each considered as its own cluster. Step 2. Define cluster distances (average linkage): The distance between two clusters ***C***_***p***_ and ***C***_***q***_ is defined as the average of all pairwise distances between points in ***C***_***p***_ and points in ***C***_***q***_:

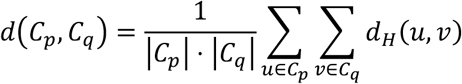

where |***C***_***p***_| denotes the number of reads in cluster ***C***_***p***_.

Step 3. Compute all inter-cluster distances: At this stage, compute the complete distance matrix between every pair of clusters.

Step 4. Merge closest clusters: Identify the pair of clusters (*C*_*r*_, *C*_*s*_) such that

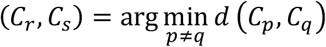

Merge these two clusters into a new cluster *C*_rs_ = *C*_*r*_ ∪ *C*_*s*_

Step 5. Update the distance matrix: Remove the rows and columns corresponding to *C*_*r*_ and *C*_*s*_. Add a row/column for the new cluster *C*_rs_. Compute distances between *C*_rs_ and each remaining cluster *C*_k_ using the average linkage formula.

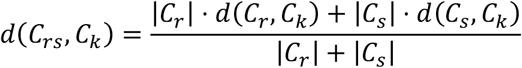

Step 6. Repeat until target cluster number is reached: Continue iteratively merging clusters until the number of clusters equals the predefined parameter *t*.

#### Cluster phasing/stitching

After per-window clustering, each of the *M* windows yields *t* clusters. To generate *t* consistent clusters across the entire region, clusters must be stitched between successive windows. This task is an assignment problem: matching clusters in window *N* to the most similar clusters in window *N* + 1. The problem can be expressed as a mathematical optimization:

Let the windows *W*_*N*_ and *W*_*N*+1_, two successive windows each containing *t* clusters

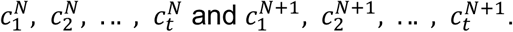

Given a distance matrix *D* = [*d*_*ij*_] with *i, j* = 1, . . ., *n* where *d*_*ij*_ is the distance between cluster 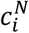 and 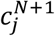, the assignment problem is to find a binary matrix **X** = [*x*_*ij*_] where each entry *x*_*ij*_ indicates whether cluster 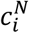 is matched to cluster 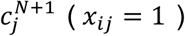 or not (*x*_*ij*_ = 0) and which minimizes the total cost:

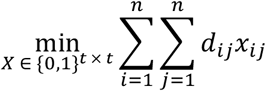

Subject to:

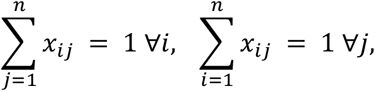

so that every cluster in window *N* is matched uniquely to one cluster in window *N* + 1

To solve this problem, we used the Hungarian algorithm, a classical method for optimal assignment in combinatorial optimization, with as input the distance matrix

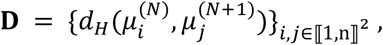

where *d*_*H*_(*u, v*) is the Masked Hamming distance between two vectors *u* and *v* and 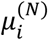 is the mean methylation sequence vector of cluster 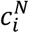

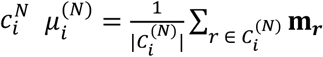, where **m**_**r**_ is the methylation sequence vector of read *r*

and the following procedure:

Step 1. Row reduction: For each row in the cost matrix **D**, subtract the minimum value of that row from all elements in the row.

Step 2. Column reduction: For each column, subtract the minimum value of that column from all elements in the column.

Step 3. Cover zeros: Cover all zeros with the minimum number of horizontal/vertical lines. If the number of lines = *t*, proceed to Step 5, otherwise, continue to Step 4.

Step 4. Adjust uncovered elements: Find the smallest uncovered value δ (i.e., not crossed by any line), subtract δ from all uncovered elements, add δ to elements covered twice (i.e., by both a row and column) then repeat Step 3.

Step 5. Find an optimal assignment: Find a zero in the matrix, assign it, and eliminate its row and column from further assignment. Continue until every row has a unique assignment. This gives you the optimal assignment minimizing total cost.

Repeating the procedure for each pair of adjacent windows yields an optimal one-to-one matching of clusters across all *M* windows. This ensures that each of the *t* stitched clusters is continuous and consistent throughout the region.

#### Global Clustering

Each read was assigned to its most similar stitched cluster series based on masked Hamming distance. Final clusters thus represented groups of single-molecule methylation profiles spanning the entire locus. Cluster assignments were assessed by plotting individual reads alongside their corresponding series profiles at the bin and GpC level.

#### Long read accuracy

To evaluate the accuracy of our clustering approach, we selected 100 loci each covered by at least one long read (>50 kb). After filtering for informative reads, 90 loci with sufficient coverage were retained (**Table S1**). For each locus, the cluster-based stitching procedure was performed while leaving out the long read. After clustering, the long read was assigned to its best-matched global cluster. For each locus, the methylation sequence of the long read was compared to the mean methylation sequence of each cluster (*c*_*i*_). The closest cluster was identified using the masked Hamming distance *d*_*H*_.

The read accuracy for a long read *r*_*i*_ was defined as:

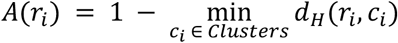

The overall clustering accuracy was defined as the average accuracy across all long reads:

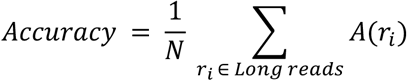

## Data availability

No data were generated in this study. Accession numbers of the datasets used are reported in the datasets section.

## Code availability

The codes for the clustered-based phasing strategy will be made publicly available on zenodo upon acceptance (https://doi.org/10.5281/zenodo.17064564).

## Acknowledgements

This work was supported by 1R35GM122515 (J.S) and NIH P01CA229086 (J.S).

The authors thank Skok lab members for helpful scientific discussions, New York University School of Medicine High Performance Computing Facility (HPCF) for computing technical support, Adriana Heguy and the Genome Technology Center (GTC) core for sequencing efforts. GTC is a shared resource partially supported by the Cancer Center Support Grant P30CA016087 at the Laura and Isaac Perlmutter Cancer Center.

## Author contributions

These studies were designed by Jane Skok, Catherine Do and Stephanie Bellini. The analysis was performed by Stephanie Bellini and Catherine Do. The paper was written by Jane Skok and Catherine Do.

## SUPPLEMENTARY INFORMATION

**Supplemental Tables**

**Table S1. Cluster-based phasing accuracy**. Table of long reads used for the accuracy estimation. This table relates to **Figure 3**.

**Table S2. Coverage and read length median for Sox2, Hoxa and Klf1 loci**. This table relates to **Figures 4-7**.

**Supplemental Figures**

**Figure S1. Clustered-based phasing of the Nano-NOMe-seq single molecules**. Cartoon depicting the agglomerative hierarchical clustering (**A**), the Hungarian algorithm (**B**) and (**C**) the Hamming distance used in the clustered based phasing pipeline. This figure relates to **Figure 2**.

**Figure S2. Cluster-based phasing of nano-NOMe-seq data reveals CTCF-independent Sox2 transcription**. Profiles of mean methylation across all 9 clusters (excluding the fully methylated and unmethylated clusters, C1 and C9) and studied windows shown in **Figure 4A**. This figure relates to **Figure 4**.

**Figure S3. Cluster-based phasing of nano-NOMe-seq data reveals long range multiway CTCF dependent loops at the *Sox2* locus. A**. Profiles of mean methylation across all 7 clusters (excluding the fully methylated and unmethylated clusters, C1 and C7) and studied windows shown in **Figure 5A**. This figure relates to **Figure 5**.

**Figure S4. Cluster-based phasing of nano-NOMe-seq data reveals long range high-order multiway CTCF independent loops between micro-compartments at the *Klf1* locus. A**. Profiles of mean methylation across all 8 clusters (excluding the fully methylated and unmethylated clusters, C8 and C9) and studied windows shown in **Figure 7A**. This figure relates to **Figure 7**.

**Figure S1.**
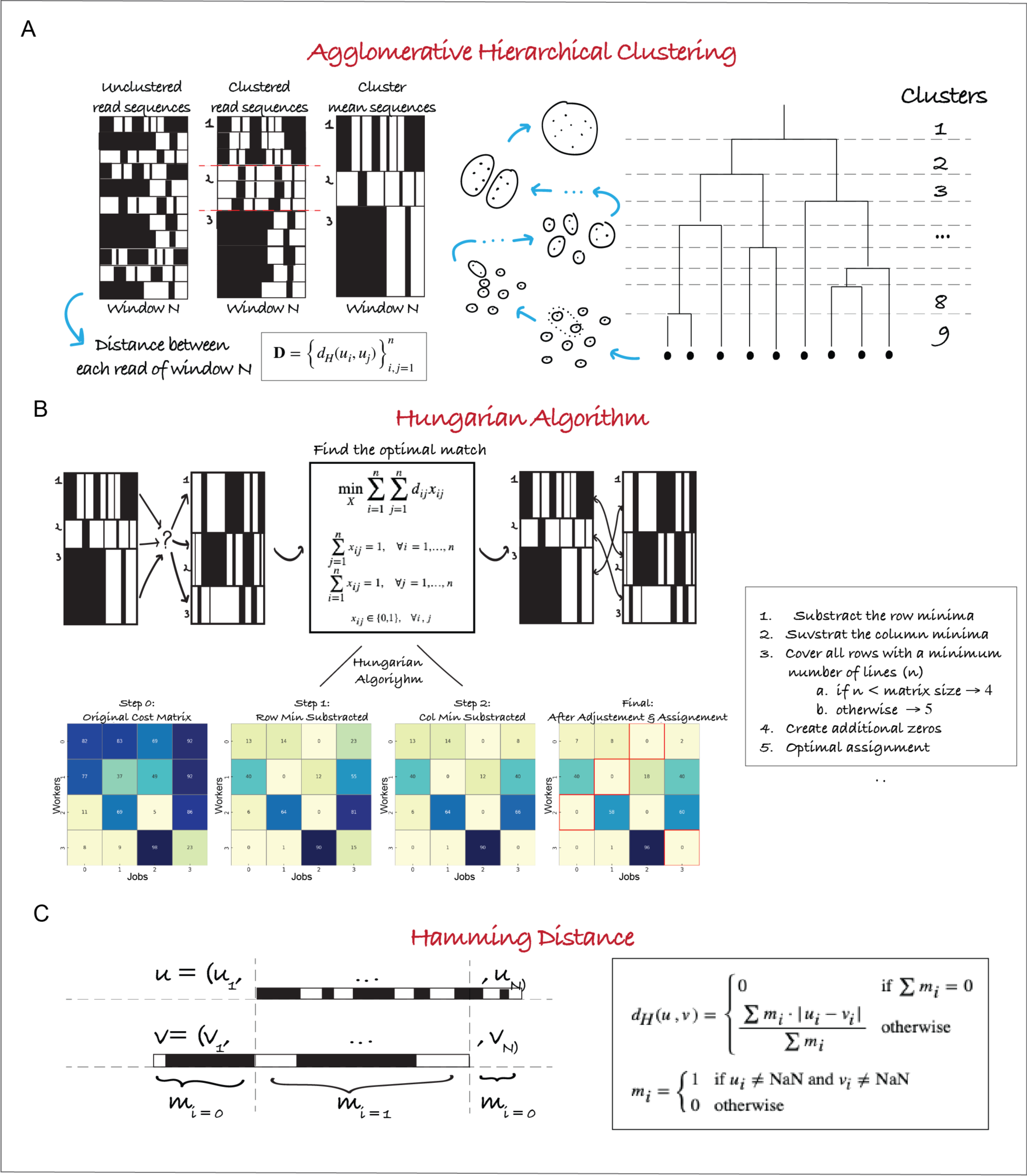
Clustered-based phasing of the Nano-NOMe-seq single molecules. Cartoon depicting the agglomerative hierarchical clustering (**A**), the Hungarian algorithm (**B**) and (**C**) the Hamming distance used in the clustered based phasing pipeline. This figure relates to **Figure 2**

**Figure S2.**
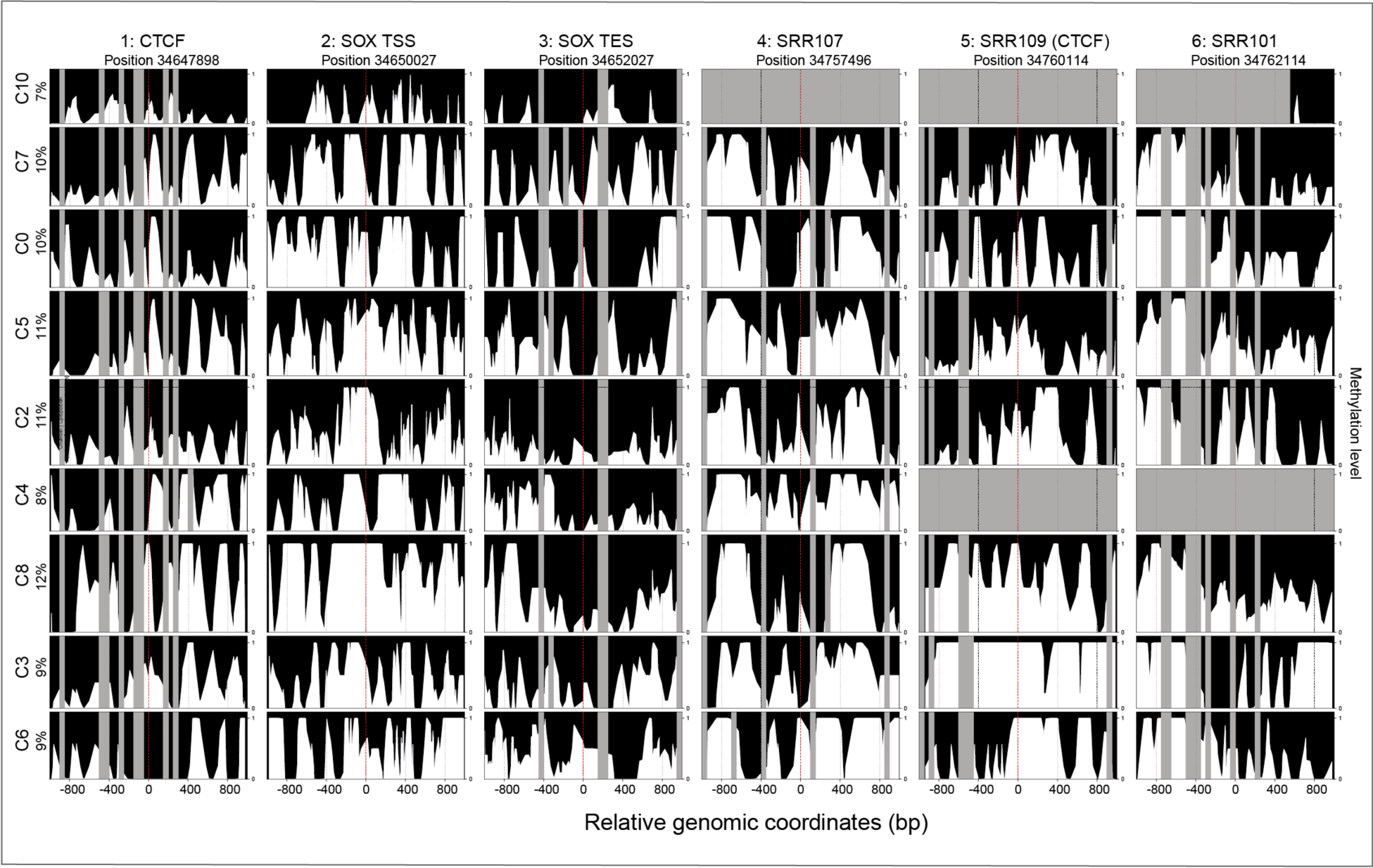
Cluster-based phasing of nano-NOMe-seq data reveals CTCF-independent Sox2 transcription. Profiles of mean methylation across all 9 clusters (excluding the fully methylated and unmethylated clusters, C1 and C9) and studied windows shown in **Figure 4A**. This figure relates to **Figure 4**

**Figure S3.**
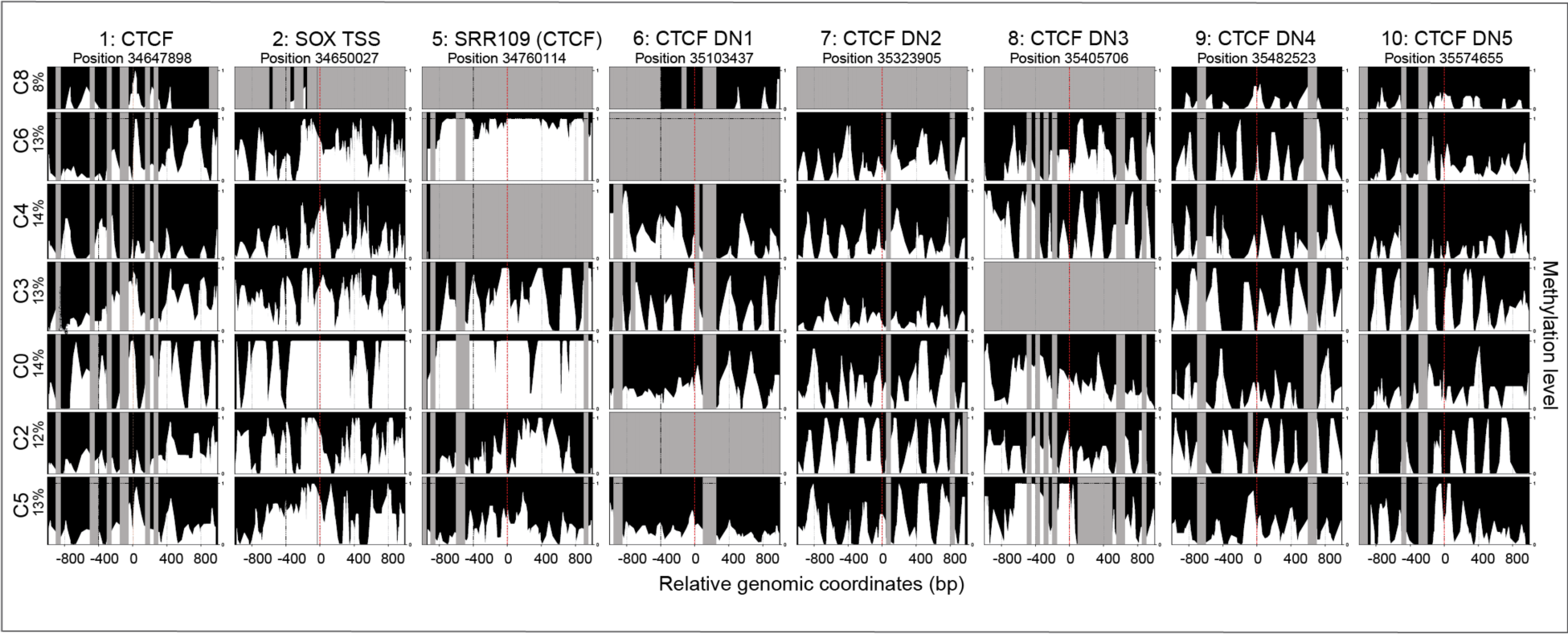
Cluster-based phasing of nano-NOMe-seq data reveals long range multiway CTCF dependent loops at the *Sox2* locus. **A**. Profiles of mean methylation across all 7 clusters (excluding the fully methylated and unmethylated clusters, C1 and C7) and studied windows shown in **Figure 5A**. This figure relates to **Figure 5**

**Figure S4.**
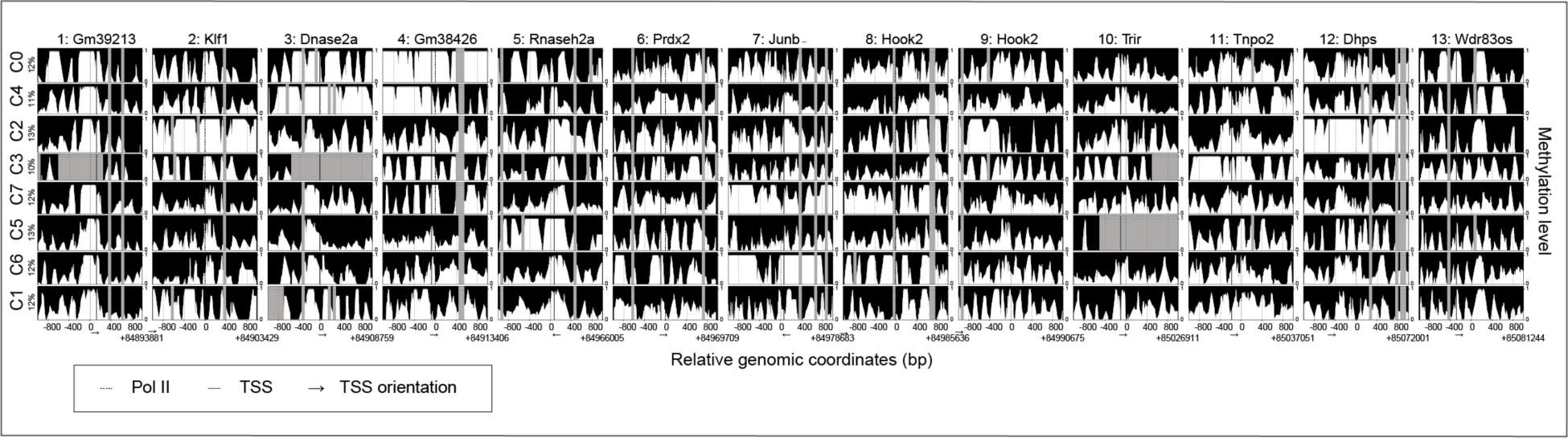
Cluster-based phasing of nano-NOMe-seq data reveals long range high-order multiway CTCF independent loops between micro-compartments at the *Klf1* locus. **A**. Profiles of mean methylation across all 8 clusters (excluding the fully methylated and unmethylated clusters, C8 and C9) and studied windows shown in **Figure 7A**. This figure relates to **Figure 7**

## REFERENCES

1. Hafner, A., Park, M., Berger, S.E., Murphy, S.E., Nora, E.P., and Boettiger, A.N. (2023). Loop stacking organizes genome folding from TADs to chromosomes. Mol Cell 83, 1377–1392 e1376. 10.1016/j.molcel.2023.04.008.

2. Ostrowski, M.S., Yang, M.G., McNally, C.P., Abdulhay, N.J., Wang, S., Renduchintala, K., Irkliyenko, I., Biran, A., Chew, B.T.L., Midha, A.D., et al. (2025). The single-molecule accessibility landscape of newly replicated mammalian chromatin. Cell 188, 237–252 e219. 10.1016/j.cell.2024.10.039.

3. Fudenberg, G., Imakaev, M., Lu, C., Goloborodko, A., Abdennur, N., and Mirny, L.A. (2016). Formation of Chromosomal Domains by Loop Extrusion. Cell Rep 15, 2038–2049. 10.1016/j.celrep.2016.04.085.

4. Dekker, J., Rippe, K., Dekker, M., and Kleckner, N. (2002). Capturing chromosome conformation. Science 295, 1306–1311.

5. Lieberman-Aiden, E., van Berkum, N.L., Williams, L., Imakaev, M., Ragoczy, T., Telling, A., Amit, I., Lajoie, B.R., Sabo, P.J., Dorschner, M.O., et al. (2009). Comprehensive mapping of long-range interactions reveals folding principles of the human genome. Science 326, 289-293. 326/5950/289 [pii]. 10.1126/science.1181369.

6. Krietenstein, N., Abraham, S., Venev, S.V., Abdennur, N., Gibcus, J., Hsieh, T.S., Parsi, K.M., Yang, L., Maehr, R., Mirny, L.A., et al. (2020). Ultrastructural Details of Mammalian Chromosome Architecture. Mol Cell 78, 554–565 e557. 10.1016/j.molcel.2020.03.003.

7. Bonev, B., and Cavalli, G. (2016). Organization and function of the 3D genome. Nat Rev Genet 17, 661–678. 10.1038/nrg.2016.112.

8. Goel, V.Y., Huseyin, M.K., and Hansen, A.S. (2023). Region Capture Micro-C reveals coalescence of enhancers and promoters into nested microcompartments. Nat Genet 55, 1048–1056. 10.1038/s41588-023-01391-1.

9. Zhang, S., Ubelmesser, N., Barbieri, M., and Papantonis, A. (2023). Enhancer-promoter contact formation requires RNAPII and antagonizes loop extrusion. Nat Genet 55, 832–840. 10.1038/s41588-023-01364-4.

10. Chang, L., Xie, Y., Taylor, B., Wang, Z., Sun, J., Armand, E.J., Mishra, S., Xu, J., Tastemel, M., Lie, A., et al. (2024). Droplet Hi-C enables scalable, single-cell profiling of chromatin architecture in heterogeneous tissues. Nat Biotechnol. 10.1038/s41587-024-02447-1.

11. Fu, Y., Rocha, P.P., Luo, V.M., Raviram, R., Deng, Y., Mazzoni, E.O., and Skok, J.A. (2016). CRISPR-dCas9 and sgRNA scaffolds enable dual-colour live imaging of satellite sequences and repeat-enriched individual loci. Nat Commun 7. 10.1038/ncomms11707.

12. Ma, H., Tu, L.C., Naseri, A., Huisman, M., Zhang, S., Grunwald, D., and Pederson, T. (2016). Multiplexed labeling of genomic loci with dCas9 and engineered sgRNAs using CRISPRainbow. Nat Biotechnol 34, 528–530. 10.1038/nbt.3526.

13. Bertrand, E., Chartrand, P., Schaefer, M., Shenoy, S.M., Singer, R.H., and Long, R.M. (1998). Localization of ASH1 mRNA particles in living yeast. Mol Cell 2, 437–445. 10.1016/s1097-2765(00)80143-4.

14. Darzacq, X., Shav-Tal, Y., de Turris, V., Brody, Y., Shenoy, S.M., Phair, R.D., and Singer, R.H. (2007). In vivo dynamics of RNA polymerase II transcription. Nat Struct Mol Biol 14, 796–806. 10.1038/nsmb1280.

15. Alexander, J.M., Guan, J., Li, B., Maliskova, L., Song, M., Shen, Y., Huang, B., Lomvardas, S., and Weiner, O.D. (2019). Live-cell imaging reveals enhancer-dependent Sox2 transcription in the absence of enhancer proximity. Elife 8. 10.7554/eLife.41769.

16. Wan, X., Kong, J., Hu, X., Liu, L., Yang, Y., Li, H., Liu, G., Niu, X., Chen, F., Zhang, D., et al. (2025). SiCLAT: simultaneous imaging of chromatin loops and active transcription in living cells. Genome Biol 26, 1. 10.1186/s13059-024-03463-9.

17. Mateo, L.J., Murphy, S.E., Hafner, A., Cinquini, I.S., Walker, C.A., and Boettiger, A.N. (2019). Visualizing DNA folding and RNA in embryos at single-cell resolution. Nature 568, 49–54. 10.1038/s41586-019-1035-4.

18. Espinola, S.M., Gotz, M., Bellec, M., Messina, O., Fiche, J.B., Houbron, C., Dejean, M., Reim, I., Cardozo Gizzi, A.M., Lagha, M., and Nollmann, M. (2021). Cis-regulatory chromatin loops arise before TADs and gene activation, and are independent of cell fate during early Drosophila development. Nat Genet 53, 477–486. 10.1038/s41588-021-00816-z.

19. Cardozo Gizzi, A.M., Cattoni, D.I., Fiche, J.B., Espinola, S.M., Gurgo, J., Messina, O., Houbron, C., Ogiyama, Y., Papadopoulos, G.L., Cavalli, G., et al. (2019). Microscopy-Based Chromosome Conformation Capture Enables Simultaneous Visualization of Genome Organization and Transcription in Intact Organisms. Mol Cell 74, 212–222 e215. 10.1016/j.molcel.2019.01.011.

20. Liu, Z., Chen, Y., Xia, Q., Liu, M., Xu, H., Chi, Y., Deng, Y., and Xing, D. (2023). Linking genome structures to functions by simultaneous single-cell Hi-C and RNA-seq. Science 380, 1070–1076. 10.1126/science.adg3797.

21. Zhou, T., Zhang, R., Jia, D., Doty, R.T., Munday, A.D., Gao, D., Xin, L., Abkowitz, J.L., Duan, Z., and Ma, J. (2024). GAGE-seq concurrently profiles multiscale 3D genome organization and gene expression in single cells. Nat Genet 56, 1701–1711. 10.1038/s41588-024-01745-3.

22. Catherine Do, G.J., Paul Zappile, Adriana Heguy, Jane A. Skok (2025). The coordination between CTCF, cohesin and TFs impacts nucleosome repositioning and chromatin insulation to define state specific 3D chromatin folding. Biorxiv.

23. Zhou, H.Y., Katsman, Y., Dhaliwal, N.K., Davidson, S., Macpherson, N.N., Sakthidevi, M., Collura, F., and Mitchell, J.A. (2014). A Sox2 distal enhancer cluster regulates embryonic stem cell differentiation potential. Genes Dev 28, 2699–2711. 10.1101/gad.248526.114.

24. de Wit, E., Vos, E.S., Holwerda, S.J., Valdes-Quezada, C., Verstegen, M.J., Teunissen, H., Splinter, E., Wijchers, P.J., Krijger, P.H., and de Laat, W. (2015). CTCF Binding Polarity Determines Chromatin Looping. Mol Cell 60, 676–684. 10.1016/j.molcel.2015.09.023.

25. Taylor, T., Sikorska, N., Shchuka, V.M., Chahar, S., Ji, C., Macpherson, N.N., Moorthy, S.D., de Kort, M.A.C., Mullany, S., Khader, N., et al. (2022). Transcriptional regulation and chromatin architecture maintenance are decoupled functions at the Sox2 locus. Genes Dev 36, 699–717. 10.1101/gad.349489.122.

26. Chakraborty, S., Kopitchinski, N., Zuo, Z., Eraso, A., Awasthi, P., Chari, R., Mitra, A., Tobias, I.C., Moorthy, S.D., Dale, R.K., et al. (2023). Enhancer-promoter interactions can bypass CTCF-mediated boundaries and contribute to phenotypic robustness. Nat Genet 55, 280–290. 10.1038/s41588-022-01295-6.

27. Do, C., Jiang, G., Cova, G., Katsifis, C.C., Narducci, D.N., Sakellaropoulos, T., Vidal, R., Lhoumaud, P., Tsirigos, A., Regis, F.F.D., et al. (2025). Binding domain mutations provide insight into CTCF’s relationship with chromatin and its contribution to gene regulation. Cell Genom 5, 100813. 10.1016/j.xgen.2025.100813.

28. Uyehara, C.M., and Apostolou, E. (2023). 3D enhancer-promoter interactions and multi-connected hubs: Organizational principles and functional roles. Cell Rep, 112068. 10.1016/j.celrep.2023.112068.

29. Hsieh, T.S., Cattoglio, C., Slobodyanyuk, E., Hansen, A.S., Darzacq, X., and Tjian, R. (2022). Enhancer-promoter interactions and transcription are largely maintained upon acute loss of CTCF, cohesin, WAPL or YY1. Nat Genet 54, 1919–1932. 10.1038/s41588-022-01223-8.

30. Core, L., and Adelman, K. (2019). Promoter-proximal pausing of RNA polymerase II: a nexus of gene regulation. Genes Dev 33, 960–982. 10.1101/gad.325142.119.

31. Deshpande, A.S., Ulahannan, N., Pendleton, M., Dai, X., Ly, L., Behr, J.M., Schwenk, S., Liao, W., Augello, M.A., Tyer, C., et al. (2022). Identifying synergistic high-order 3D chromatin conformations from genome-scale nanopore concatemer sequencing. Nat Biotechnol 40, 1488–1499. 10.1038/s41587-022-01289-z.

